# The birth-death diffusion leading to present-day Mammal diversity

**DOI:** 10.1101/2022.08.09.503355

**Authors:** Ignacio Quintero, Nicolas Lartillot, Hélène Morlon

## Abstract

Dramatic spatial, temporal and taxonomic variation in biodiversity is ultimately explained by differences in speciation and extinction rates. Mammals represent a ∼200 My old radiation that resulted in over 6500 extant species, with stark temporal, spatial and taxonomic heterogeneity in biodiversity. Throughout their history, every mammal lineage is expected to have undergone diversification rates that vary instantaneously in time resulting from the complex interplay of context-specific extrinsic factors (*e.g*., K-Pg mass extinction event, rise of angiosperms) with their evolving ecologies (*e.g*., body size, diet). When studying the diversification history of a clade, however, mathematical and computational limitations have hindered inference of such a flexible birth-death model where speciation and extinction rates evolve continuously along a phylogenetic tree. Here we overcome these challenges by implementing a series of phylogenetic models in which speciation and extinction rates are inherited and diffuse following a latent Geometric Brownian motion process. We enable full Bayesian inference using data augmentation techniques to sample from the posterior distribution of model parameters, including augmented phylogenetic trees and validate using simulations. Using a genome-informed time-calibrated tree for over 4000 Mammals species, we are able to estimate a complete and fine-grained picture of the variation in diversification rates that captures both global and lineage specific effects. We find that, contrary to the idea of a suppressed mammalian diversification before the K-Pg mass extinction event (*i.e*., explosive- or delayed-rise), mammal speciation rates dramatically increased around 10-20 My before the K-Pg. Our new model opens exciting possibilities in disentangling the drivers behind variation in diversification and assaying how small-scale processes scale-up to macroevolutionary dynamics.

## Introduction

Understanding the tempo and mode in which lineages diversify is fundamental in explaining the origin and maintenance of biodiversity. The rates at which species originate or go extinct result from the interplay between their intrinsic traits and their specific abiotic and biotic environment (Benton, 2009). For instance, environmental oscillations and landscape heterogeneity are commonly posited as major drivers of diversification by precipitating population dispersal and fragmentation as well as opening new opportunities (Barnosky, 2001; Jablonski, 2008). Similarly, the evolution of both intrinsic (*e*.*g*., species phenotype, their niche, and their evolutionary rate) and extrinsic biotic factors (*e*.*g*., competition and other inter-specific interactions) are thought to affect the pace of evolutionary radiations (Van Valen, 1973; Benton, 2009; Quintero and Landis, 2019). Therefore, the interrelation of fluctuating context-specific dynamics with species’ intrinsic evolving ecologies are expected to result in lineage-specific diversification rates that themselves evolve and can vary at any point in time.

The mammal radiation started at around 200 Mya (Upham et al., 2019; Álvarez-Carretero et al., 2021) and resulted in an estimated present-day diversity of *ca*. 6450 currently recognized species (Mammal Diversity Database, 2022). Distinct mammalian evolutionary routes led to marked differences in ecomorphologies, reflected in body size (10^8^ fold differences), generation time, litter size and habitat (aquatic, arboreal, terrestial, fossorial, etc.) variation, concomitant with an uneven distribution of richness across clades (Davies et al., 2008; Meredith et al., 2011; Grossnickle et al., 2019). Furthermore, throughout their long evolutionary history, lineages were differentially impacted by environmental factors such as the radiation of flowering plants (*i*.*e*., the Cretaceous Terrestrial Revolution), the K-Pg extinction event, the Paleocene-Eocene thermal maximum, and other dramatic environmental oscillations, which likely spurred widespread distribution shifts and extinctions together with novel ecological opportunities that impacted diversification rates (Meredith et al., 2011; Grossnickle et al., 2019; Upham et al., 2021). Indeed, a major unsolved debate in mammalian evolution revolves around understanding the timing at which the orders with living representatives originated and diversified (Bininda-Emonds et al., 2007; Stadler, 2011a; Meredith et al., 2011; Grossnickle et al., 2019; Springer et al., 2019; Upham et al., 2021). Standing hypotheses posit that crown orders increased their diversification either before, at or after the K-Pg event, dubbed as ‘early-’, ‘explosive-’ or ‘delayed-’ rise of extant mammals, respectively (Meredith et al., 2011; Grossnickle et al., 2019). The complex interplay of mammal species-specific ecomorphologies with their particular environments that fluctuate throughout lineage’s duration translate into lineage- and time-specific rates of diversification along their evolutionary history, that is, a given lineage is expected to undergo diversification rate changes at any moment in time. Nonetheless, extreme external events, such as the rise of angiosperms or the K-Pg extinction event, are thought to transcend lineage-specific diversification dynamics, and leave a common signature across lineages during that particular period (Barnosky, 2001). Therefore, to explore the temporal dynamics that led to extant mammal diversity, an idealized model of diversification that enable the reconstruction of overarching temporal dynamics while incorporating rates of speciation and extinction that change instantaneously along time for any lineage is needed.

Several phylogenetic approaches have been developed to account for heterogeneity in time or across lineages in characterizing diversification rate variation across taxa. Most available methods that incorporate rate heterogeneity across lineages assume a (usually few) number of independent shifts across the tree that partition the tree into separate ‘regimes’ (or states) wherein lineages undergo constant rates: ‘BAMM’ (Rabosky, 2014), ‘birth-death-shift’ process (Höhna et al., 2019), ‘MTBD’ (Barido-Sottani et al., 2020), and ‘MiSSE’ (Vasconcelos et al., 2022). Instead of identifying large diversification shifts along a phylogenetic tree, ‘ClaDS’ assumes that shifts occur at each cladogenetic event, with daughter lineages undergoing rate constancy within each lineage (Maliet et al., 2019). The only method we are aware of that allows lineage-specific speciation and extinction rates to vary instantaneously through time, ‘QuaSSE’, assumes that this variation is completely explained by the evolution of a trait under Brownian motion (FitzJohn, 2010). While these methods have proven very useful in reconstructing diversification dynamics on phylogenetic trees, they remain restrictive in assuming rate constancy either across ‘regimes’ or along a given branch, and a more flexible model is required.

Here we assess the tempo and mode in which surviving mammal orders originated and diversified by applying the first phylogenetic diversification model in which lineage-specific diversification rates vary instantaneously in time by assuming that speciation and extinction rates follow a Geometric Brownian motion: the Birth-Death Diffusion (BDD) model. We show that the BDD and other simpler diffusion models, including no extinction, constant-extinction and constant-turnover, exhibit good statistical properties in most scenarios and are identifiable even when restricted to extant taxa alone. Considering that the temporal dynamics of mammalian diversification will rely on the accuracy of the phylogenetic tree, we use the latest time-calibrated molecular tree for mammals, which incorporated 72 species genomes enhancing the time-dating robustness (Álvarez-Carretero et al., 2021). We validate our method and perform Bayesian inference of our diffusion models on this mammalian tree using data augmentation techniques. On top of posterior probabilities for the main process parameters, our model also returns for free posterior sampled histories of diversification across trees with unobserved speciation events enabling post-hoc analyses and visualizations.

## Model

We assume that, at some time *t*, each lineage *l* has an instantaneous rate of producing new species of λ_*l*_(*t*) (*i*.*e*., speciation rate) and an instantaneous rate of going extinct of *µ*_*l*_(*t*) (*i*.*e*., extinction rate). This general birth-death process generates a bifurcating phylogenetic tree with some lineages dying out and others giving rise to daughter species after some time. Probabilistic inference is complicated since, in practice, we do not observe the whole process, but rather the evolutionary relationships among those lineages that were able to be sampled, that is, the “reconstructed” phylogenetic tree (Nee et al., 1994). Our goal is to perform inference on speciation and extinction rates that are inherited and stochastically diffuse through time following a Geometric Brownian motion, given that we only observe the reconstructed tree. Because there is no available analytical solution to estimate the likelihood, we use Bayesian data augmentation techniques to perform full posterior inference on birth-death diffusion models. We start by describing the data augmented approach for a simple constant rate birth-death (‘CBD’) model, which we then expand on to enable inference on models with rate diffusion.

Let Ψ be an ultrametric rooted phylogenetic tree under a general birth-death process that starts at some time *T*_Ψ_ (*i*.*e*., tree height) in the past and continues to time 0 in the present with *n* surviving tips and per-lineage speciation rate λ_*l*_(*t*) and extinction-rate *µ*_*l*_(*t*) for lineage *l* at time *t*. Furthermore, each clade *c* has a specific probability for extant lineages to be represented in the observed tree, specified by *ρ*_*c*_ ∈ [0, 1]. We propagate this probability throughout all the branches in the tree by specifying a branch-specific sampling fraction *ρ*_*b*_ for branch *b*. For terminal branches in clade *c* we simply assign *ρ*_*b*_ = *ρ*_*c*_ ∀*b* ∈ *c*. Let *a*_*b*_ be the number of alive tips descending from branch *b* in the observed tree, then, for internal branches, we calculate *ρ*_*b*_ = (*a*_*d*1_ + *a*_*d*2_)/(*a*_*d*1_/*ρ*_*d*1_ + *a*_*d*2_/*ρ*_*d*2_), where *d*_1_ and *d*_1_ are the daughter branches. Thus, the probability of sampling exactly 1 extant species for a terminal branch *b* is 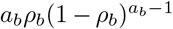 and sampling no extant species for internal branch *b* is (1 − *ρ*_*b*_)^*a*− *b*^, following Maliet and Morlon (2021).

We use Bayesian data augmentation (DA) to sample unobserved lineages that either went extinct in the past or were not sampled at the present during inference. For clarity let Ψ_*o*_ represent only the reconstructed (observed) tree, Ψ_*a*_ represent some unobserved speciation and extinction events and let Ψ (= Ψ_*o*_ ∪ Ψ_*a*_) represent the complete tree. Given that we do not observe Ψ_*a*_, we treat it as a random variable to integrate over using Markov Chain Monte Carlo (MCMC).

### Constant rate Birth-Death

In the CBD model, at any time, all lineages share the same speciation and extinction rate λ and *µ*, respectively (*i*.*e*., λ_*l*_(*t*) = λ and *µ*_*l*_(*t*) = *µ*).

#### Likelihood

The likelihood for a complete unordered crown tree under a CBD process is simply

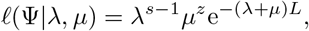

where *s* is the number of speciation events, *z* is the number of extinction events, *L* is the tree length (sum of all branches). For a stem tree, we do consider all *s* speciation events.

To integrate over Ψ during MCMC, we developed two alternative approaches for data augmentation. A first approach samples Ψ_*a*_ directly from its conditional distribution, given Ψ_*o*_ and the model parameters (*i*.*e*., Gibbs sampling) using forward simulation. A second approach is based on accept-reject sampling, based on proposals of grafting and pruning. Each can be useful on different contexts, but we relied mostly on the forward simulation approach, which we describe below, and leave the description of the grafting/pruning approach for the Appendix.

#### Forward simulation

Here we follow Maliet and Morlon (2021) in our forward simulation approach but describe it more generally for the CBD process. First, we uniformly sample a branch in the tree and simulate a birth-death process forward in time throughout the branch length *t*_*b*_ for branch *b*, given the current model parameters. For any branch, if the process becomes extinct before *t*_*b*_, we reject the proposal (*i*.*e*., it requires that at least 1 species is alive at time *t*_*b*_). Note that for a terminal branch *t*_*b*_ is the present. If the branch is internal, and there is more than one surviving lineage by *t*_*b*_, we pick one tip at random as the one that gave rise to the observed speciation event and continue the simulation in forward fashion for the others. All other unobserved lineages should go extinct before the present or, if there are unobserved data augmented lineages at the present, then the proposal is rejected or accepted following the branch’s sampling fraction, *ρ*_*b*_.

Let *ψ*_*b*_ denote the forward simulated tree in branch *b, n*_*b*_(*t*_*b*_) denote the number of lineages alive at time *t*_*b*_ (*i*.*e*., at the end of the branch) and *n*_*b*_(0) denote the number of lineages alive at the present for *ψ*_*b*_, then the acceptance ratio in the Metropolis-Hastings (MH) step for the proposed forward simulation 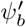 if *b* is a terminal branch is

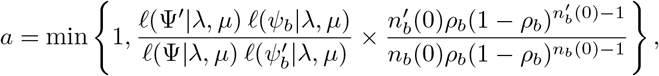

and if *b* is an internal branch

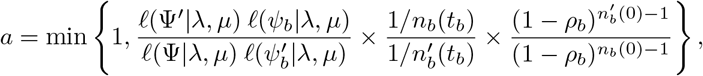

where the 1/*n*_*b*_(*t*_*b*_) factor comes from the proposal density of randomly and uniformly choosing one of the extant lineages at time *t*_*b*_ as the one sampled in the reconstructed tree. Because the full tree likelihood for the proposal only differs on the specific branch, the acceptance ratio simplifies in terminal branches to

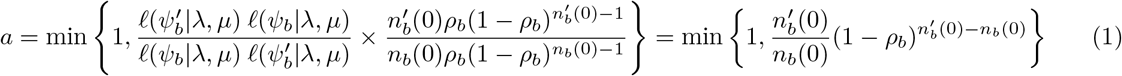

and in internal branches to

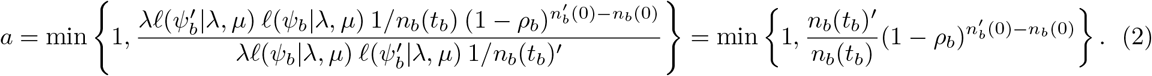

The acceptance probability for the sampling fraction is common to all further models, so, for simplicity, we suppress it in the rest of the manuscript.

#### Conditioning on survival

To condition on survival of the process we use the pseudo-marginal principle from Andrieu and Roberts (2009). We describe here the approach in a general manner since it applies to all models that involve extinction. Assuming crown conditioning, that is, that both crown lineages survive to the present and letting *f*(Ψ|*θ*) be the likelihood of the tree and *p*(*θ*) the prior for parameters *θ* we target the unnormalized density

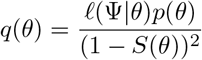

where *S*(*θ*) is the probability that a lineage at *T*_Ψ_ survives until the present.

Following Ronquist et al. (2021),

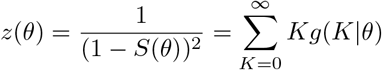

where *g*(*K*|*θ*) = (1 − *S*(*θ*)^2^)^*K*−1^*S*(*θ*)^2^. Thus, *z*(*θ*) is the expectation of a geometric distribution of parameter *S*(*θ*)^2^. We can then sample *K* from *g*(*K*|*θ*) (for which we do not have an analytical solution: see Appendix), by simulating two lineages under *θ* starting at *T*_Ψ_ and counting the number of attempts until both survive.

To sample from density *q*(*θ*) we perform MCMC on the pair of variables (*θ, K*) targeting instead

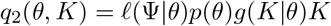

marginally obtaining our target density

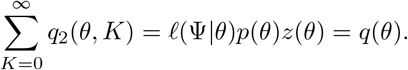

Effectively, our algorithm proposes a new value for *θ, θ*′ ∼ *s*(*θ, dθ′*), and then, given *θ*′, proposes *K* by drawing from *K* ∼ *g*(*K* |*θ*′) and accept this with probability *a* = min{1, *r*}, where

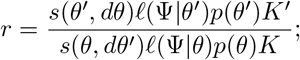

circumventing the need to analytically compute *z*(*θ*) while providing exact MCMC inference. Of note, in this equation for r, ℓ(Ψ|*θ*)*p*(*θ*)*K* is an unbiased estimate of *q*(*θ*) in the denominator, and similarly for *q*(*θ*′) in the numerator. Replacing the target by an unbiased stochastic estimate of it in the acceptance ratio, while preserving exact MCMC sampling, is the fundamental idea of the pseudo-marginal principle (Andrieu and Roberts, 2009). For stem conditioning, the probability of survival of one lineage until the present, we simply estimate *K* as the number of times until a single lineage at time *T*_Ψ_ survives to the present (*i*.*e*., *z*(*θ*) is the expectation of a geometric distribution of parameter *S*(*θ*)).

#### Mixed Gibbs sampling for λ and *µ*

Given the complete tree, Ψ, λ and *µ* follow a Poisson distribution and thus one can sample (almost) directly from their full conditional posterior via Gibbs sampling. We use the conjugate Gamma prior, Γ(*κ, ς*) for both λ and *µ*, which results in the following full conditional distribution from which we can sample directly

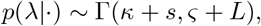

and similarly with *µ* but using the number of extinction events. The conditioning on survival, however, requires adding a MH step, such that the proposed speciation and extinction rates λ′ and *µ*′ are accepted with probability *a* = min{1, *K*′ /*K*}.

We validate our data augmentation implementation of the constant rate birth-death by comparing the rate posteriors with the analytical solution from (Nee et al., 1994) implemented in a Bayesian framework in the ‘diversitree’ package (FitzJohn, 2010) for R (R Core Team, 2022) (see Fig S1).

### Pure-Birth Diffusion

We now relax the condition that speciation rates are constant through time and across taxa by rather defining λ_*l*_(*t*) to be the result of a stochastic diffusion process. In other words, we consider the observed phylogenetic tree as the outcome of an unobserved latent process of speciation. For simplicity, we start by assuming that there is no extinction, and call this model the Pure-Birth Diffusion (‘PBD’). Specifically, we assume that speciation rates for lineage *l* evolve anagenetically following the exponential of a Brownian motion (*i*.*e*., Geometric Brownian motion, GBM), such that

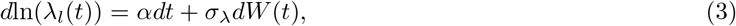

where *α* represents the drift and *σ*_λ_ the diffusion rate for speciation rates and *W* (*t*) is the Wiener process (*i*.*e*., standard BM). This diffusion rate has units 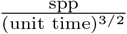. Under this model, the drift, *α*, determines the median or geometric expectation of the rates, but not its arithmetic mean expectation. That is, if *α* = 0, after some time half of the GBM processes will be lower than the starting rate and the other half will have increased. Given the non-negative nature and log-normal expectation, the arithmetic mean of will be larger than the median, which is why we perform all rate transformations and aggregations using the geometric mean (see below).

At cladogenesis, the current value for speciation rates are inherited identically, that is, for any cladogenetic event at time *t*_*s*_ and an ancestral lineage rate of λ_*a*_(*t*_*s*_), then λ_*d*1_ (*t*_*s*_) = λ_*d*2_ (*t*_*s*_) = λ_*a*_(*t*_*s*_), where *d*_1_ and *d*_2_ represent each of the daughter lineages. At the start of the tree (*i*.*e*., time *T*_Ψ_) λ_*i*_(*T*_Ψ_) = λ_*r*_.

Note that adding the drift parameter *α* is important in some contexts to restrain the “run-away species selection” that our model and others with inherited speciation rates (Beaulieu and O’Meara, 2015; Maliet et al., 2019), produce. Specifically, lineages with higher speciation rates will generate daughters with a higher expected rate, which will themselves generate even more daughters, and so on. Including a drift parameter and/or extinction constraints allows avoiding such a run-away.

#### Data augmentation

We use a data augmentation approach to approximate the likelihood of rates that follow GBM diffusion (Horvilleur and Lartillot, 2014; Quintero and Landis, 2019). Namely, we generate unobserved stochastic paths of GBM for all lineages across the tree to approximate the likelihood of a phylogenetic tree generated under Eq. 3. We determine a small time step, *δ*, such that *δ*_*i*_ = *t*_*i*+1_ − *t*_*i*_, where *t*_*i*_ < *t*_*i*+1_, and divide each branch of the tree into *m* small time steps such that *m* = *t*_*b*_*/δ* + 1, where *t*_*b*_ represents the edge length of branch *b*. Note that the last time step for branch *b* is always smaller than *δ* assuming that the probability that *t*_*b*_ is a multiple of *δt* is 0. Thus, for any branch *b*, we sample the data augmented diffusion process at times *t* = {*t*_1_ = 0, *t*_2_ = *t*_1_ +*δ*, …, *t*_*m*+1_ = *t*_*b*_}, obtaining the stochastic process Λ_*b*_ = {λ_*b*_(*t*_1_), …, λ_*b*_(*t*_*b*_)} sampled in a discrete time grid. For clarity and conciseness we denote λ_*l*_(*t*_*i*_) as λ_*i*_ from now on.

#### Likelihood

For any time step [*t*_*i*_, *t*_*i*+1_], we sample λ_*i*_ and λ_*i*+1_ at the endpoints. Given a sufficiently small time step, *δ*_*i*_, a good approximation for the likelihood of no event happening during time [*t*_*i*_, *t*_*i*+1_] is:

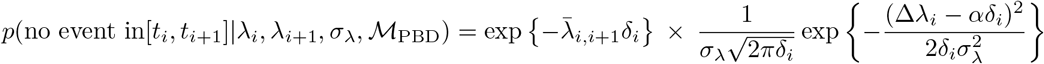

where 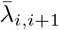 represents the geometric mean for {λ_*i*_, λ_*i*+1_}, Δλ_*i*_ = ln(λ_*i*+1_) − ln(λ(*t*_*i*_)), and ℳ_PBD_ represents the PBD model given by Eq. 3. The first part of the equation is the probability that there is no speciation events during *δ*_*i*_ and the second part is the probability that the GBM diffused from λ_*i*_ to λ_*i*+1_. For internal branches we then simply multiply the speciation events likelihoods. Given the data augmented stochastic diffusion for speciation rates across the tree, Λ, the likelihood for the full tree under the Yule Diffusion process can be then straightforwardly approximated by

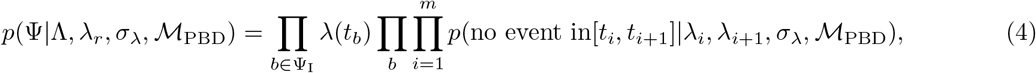

where Ψ_I_ are the set of internal branches. Note that topology matters, and for an ancestral lineage *a* with daughter branches *d*_1_ and *d*_2_, we have that 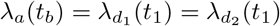

#### Diffusion updates

We integrate over possible rate diffusion histories, Λ, using Metropolis-Hastings diffusion path updates. We use a path diffusion update based on internal nodes that are connected to a ‘triad’ of branches in Ψ, that is, given a parent and its daughter branches, *pr, d*_1_ and *d*_2_, respectively. We consider the following four cases to update the triad diffusion paths Λ_*pr,d*1_,*d*_2_: *i)* If all branches are internal and the parent is not the root, we make a GBM proposal for the node conditioned on the end points (i.e., λ_*pr*_(*t*_1_), λ_*d*1_ (*t*_*b*_), λ_*d*2_ (*t*_*b*_)) and respective branch lengths. We then propose diffusion paths using Brownian bridges for each branch with the new node value as an endpoint. *ii)* If all branches are internal and the parent is the root, we make a GBM node proposal conditioned on the daughter end points, from which we then backwardly propose a new root value λ_*r*_. We then use Brownian bridges to sample their respective diffusion paths. *iii)* If one of the daughter branches is terminal, we make a GBM node proposal conditioned on the endpoints, from which we propose a new GBM value for the endpoint (*i*.*e*., tip) of the terminal branch. We then use Brownian bridges to sample their respective diffusion paths. *iv)* If both of the daughter branches are terminal, we simply make a GBM node proposal given the parent branch, and then forwardly simulate both terminal daughter diffusion paths. Note that the drift term, *α*, cancels out when proposing Brownian bridges with drift, as it is fully determined by the endpoints and time elapsed.

Then, the acceptance probability, *a*, for a new diffusion path proposal, Λ ′ is

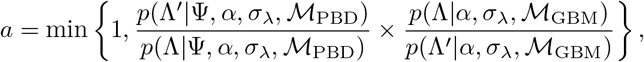

where ℳ_GBM_ represents the GBM model. Note that this ratio simplifies because the Brownian motion part in the likelihood cancels out with the proposal probability. We specify a uniform prior on the the speciation rates at the root, λ_*r*_.

#### *α* and *σ*_λ_ updates

We use a Normal conjugate prior, *p*(*α*) ∼ N(*ν, τ*) to directly sample from the full conditional posterior distribution (see Appendix for derivation):

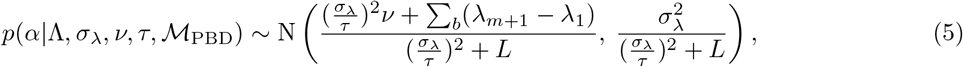

where *L* is the tree length. Similarly, we specify the Inverse Gamma conjugate prior 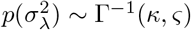, and we obtain the following posterior conditional distribution (see Appendix for derivation):

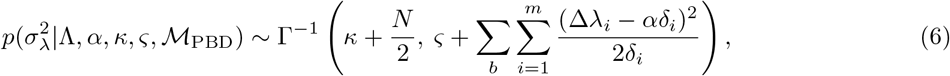

where 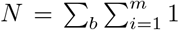.

While the Inverse Gamma prior for *σ*_λ_ is advantageous by allowing full Gibbs sampling, substantially enhancing the properties of our MCMC approach, it has some drawbacks, particularly with small phylogenetic trees. As expected, small datasets (*e*.*g*., 50 tip trees) are particularly affected by the choice of prior: a Γ^−1^(0.05, 0.05), for instance, will enforce very low probabilities to small values (*e*.*g*., *σ*_λ_ < 0.1), overestimating the diffusion rate. Conversely, a Γ^−1^(0.1, 0.001) prior, for instance, will capture small values of *σ*_λ_ but will underestimate higher diffusion rates (Fig S2). While the prior choice becomes asymptotically irrelevant as more data is included, we emphasize that, for inference, some rationale should be taken for the choice of prior (Fig S2).

### Birth-Death Diffusion

We now relax the condition that extinction rates are non-existent and rather define three diffusion models that incorporate extinction: *i)* constant extinction, *µ*(*t*) = *µ* (‘CED’), *ii)* constant turnover, *µ*(*t*) = *Eλ*(*t*) (‘CTD’), and *iii)* extinction also being the result of a a stochastic diffusion process, *µ*(*t*), (‘BDD’). For conciseness, we now describe only the latter, the Birth-Death Diffusion (BDD), which is the most general among these models, with straightforward simplifications towards the CED and CTD. Thus, we assume that extinction rates for lineage *l* evolve anagenetically following the exponential of a Brownian Motion (i.e., Geometric Brownian motion, GBM) with no drift, such that

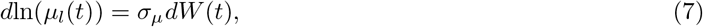

where *σ*_*µ*_ represents the diffusion rate for extinction rates and *W* (*t*) is the Wiener process (i.e., standard BM). At cladogenesis, the current value for extinction rates is inherited identically, in the same manner as are speciation rates. At the start of the tree, with time *T*_Ψ_, λ_*l*_(*T*_Ψ_) = λ_*r*_ and *µ*_*l*_(*T*_Ψ_) = *µ*_*r*_. Thus, the model has three hyper-parameters: *α, σ*_λ_, and *σ*_*µ*_. As we did for λ_*i*_, we denote *µ*_*l*_(*t*_*i*_) as *µ*_*i*_ for conciseness.

#### Likelihood

In the same way of the PBD process, for any time step [*t*_*i*_, *t*_*i*+1_], we sample λ_*i*_, λ_*i*+1_, *µ*_*i*_ and *µ*_*i*+1_ at the endpoints. Given a sufficiently small time step, *δ*_*i*_, a good approximation for the likelihood of no events during time [*t*_*i*_, *t*_*i*+1_] is:

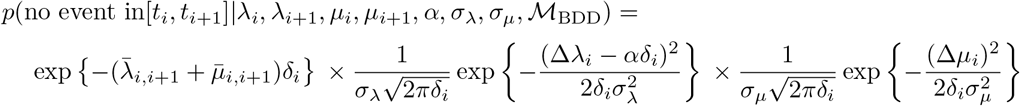

where and ℳ_BDD_ represents the Birth-Death Diffusion model given by Eq. 7. The first part of the equation is the probability that no speciation or extinction events happen during *δ*_*i*_ and the second part is the probability of the speciation and extinction GBM diffusing from λ_*i*_ to λ_*i*+1_ and *µ*_*i*_ to *µ*_*i*+1_, respectively. Given the data augmented stochastic diffusion for speciation and extinction rates across the tree, Λ and M, respectively, the likelihood for the full tree under the BDD process can be then straightforwardly approximated by

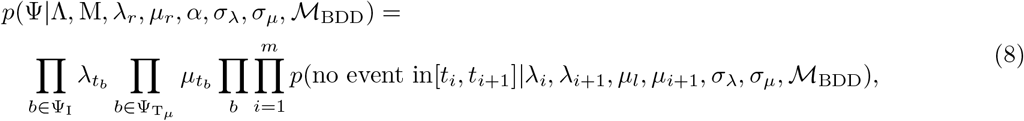

where 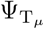 are the set of terminal extinct branches.

#### Data augmentation and parameter updates

To augment the reconstructed tree and obtain complete trees, we use forward simulation, as described above. For internal branches, new complications arise given the underlying GBM in speciation and extinction. Specifically, once the tree is simulated along a branch and one of the tips selected at random as the one that leads to the observed speciation event, both the speciation rate instantaneously before speciation λ(*t*_*pr*_) ′ and the extinction rate *µ*(*t*_*pr*_)′ do not correspond to the initial speciation and extinction rate in the daughter lineages, λ_*d*1_(*t*_1_) and λ_*d*2_(*t*_1_) and *µ*_*d*1_(*t*_1_) and *µ*_*d*2_(*t*_1_). Thus, we make Brownian bridge proposals on the daughter branches to match the new proposed rates at speciation, λ(*t*_*pr*_)′ and *µ*(*t*_*pr*_)^′^. Figure 1 illustrates our forward simulation proposals. Let Ψ, Λ, M represent the topology, the latent speciation GBM and the latent extinction GBM, respectively, then the acceptance ratio for this proposal is

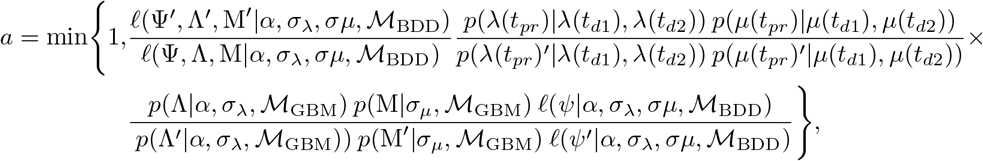

which simplifies to

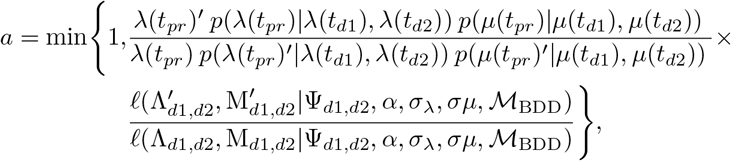

where

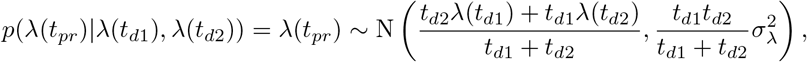

and similarly for *p*(*µ*(*t*_*pr*_)|*µ*(*t*_*d*1_), *µ*(*t*_*d*2_)). If the branch is internal we add the factor *n*_*b*_(*t*_*b*_) */n*_*b*_(*t*_*b*_) as in Eq. 2. In practice, to make proposals more efficient, we sample the tip that leads to the observed speciation event proportional to the probability that it’s rates would yield the currently observed rates at the daughters.

**Figure 1.**
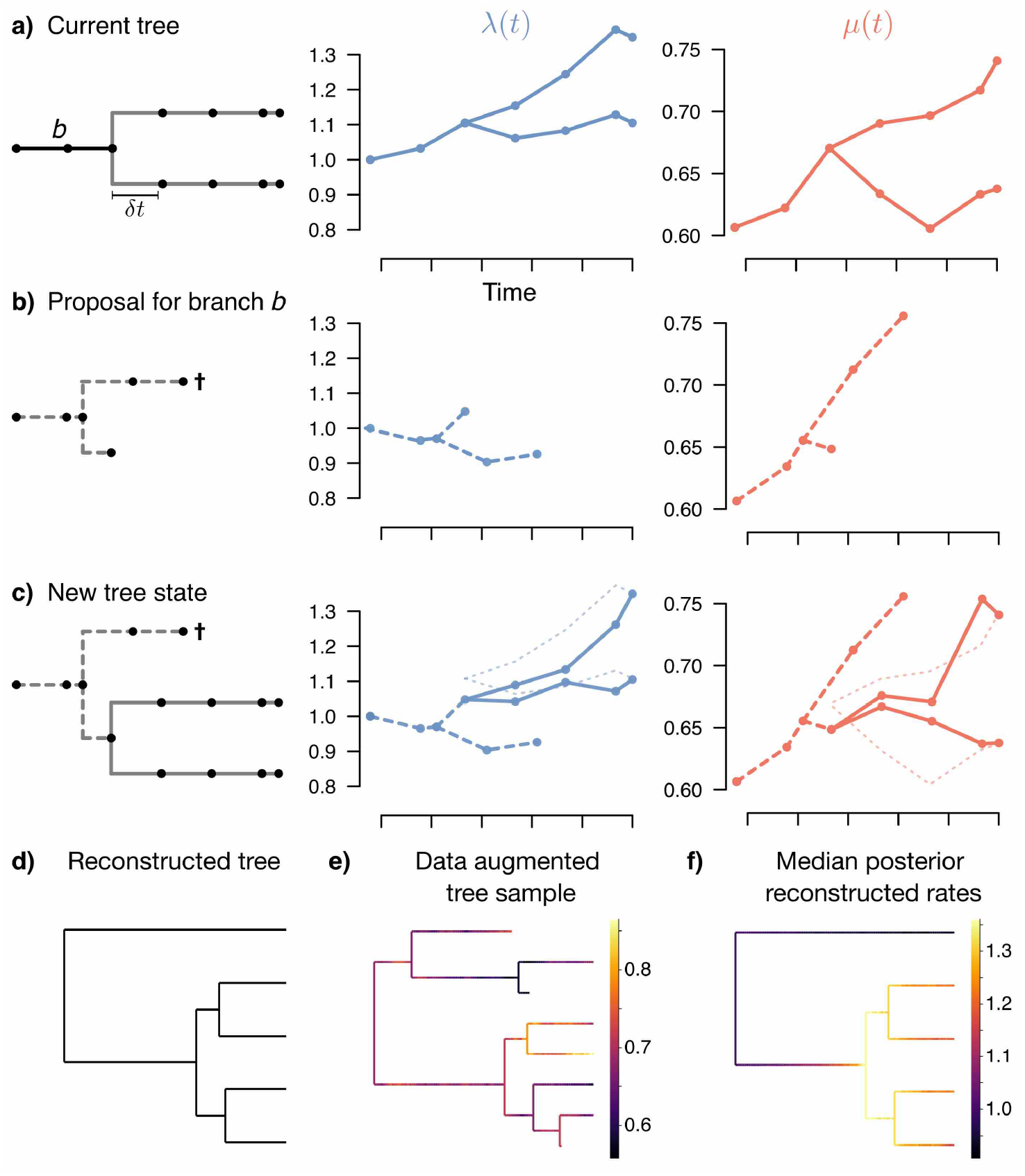
Tree BDD data augmentation. **a)** A phylogenetic tree is divided by a discrete time grid every *δ* unit times (or less) with underlying speciation λ(*t*) (blue) and extinction *µ*(*t*) rates (red). **b)** To make a new subtree proposal for branch *b*, we simulate the process with the current configuration of parameters for the duration of the branch. Here, two tips were simulated for this duration, so we picked one at random to be the one leading to the observed speciation event and we continue the simulation for the rest (here the other tip went extinct). **c)** Since the new proposal in branch *b* has different speciation and extinction rates at the tip than the current subtree of branch *b* (λ(*t*) and *µ*(*t*) thin dashed lines), we also propose rates in the daughters that match the proposal rates configuration (λ(*t*) and *µ*(*t*) solid lines), and accept or reject according to the MH acceptance ratio. **d)** An example reconstructed tree. **e)** A given data augmented MCMC sample when running the BDD method on the tree from *d)*. **f)** Median posterior reconstructed λ(*t*) from the BDD model across the tree. Color gradient represents lineage-specific variation in speciation rates along the tree, with warmer colors specifying higher rates.

We use the Eq. 5 to sample *α*, use Eq. 6 to sample *σ*_λ_, and to sample *σ*_*µ*_ we use

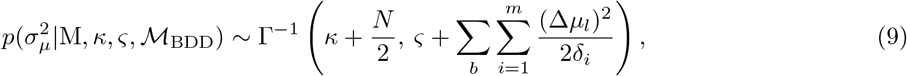

where M represents the full diffusion of *µ*(*t*) and 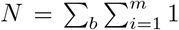. For λ_*r*_ and *µ*_*r*_ we use Uniform priors of (0, 100). We note that other sort of priors, particularly on extinction, can severely affect the posterior distribution when there is not sufficient information in the tree. For the CED and CTD models, we sample *µ* using Gibbs sampling as in the CBD model and *E* using standard MH updates and specify a Uniform prior of (0, 100).

### Model Behavior

#### Simulations

We use simulations to explore behavior for each of the four different model assumptions: PBD, CED, CTD, and BDD. To simulate under a diffusion model we take advantage of the following approximation. Let *V* be a random variable for the time of an event and let λ_*i*_ and λ_*i*+1_, be the event rate at time *t*_*i*_ and *t*_*i*+1_, respectively, where *t*_*i*+1_ − *t*_*i*_ = *δ* ≥ 0, then we have

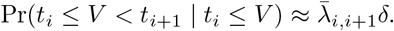

For each model we simulated 100 trees with 50, 100 and 200 tips by sampling from a range of parameter space (see Fig 2). We first made simulations and inference without *α* = 0, and then simulations with a range of values of *α* around 0 (Fig 2 & 3). To simulate non-biased samples from the model, we started with two lineages and made sure both of them survived until the present and followed Stadler (2011b) to sample a tree with a determined number of extant species. In total we have 100 simulations for each of the 4 models, each with 3 different tree sizes, with and without *α*, yielding 2400 total simulations. We ran MCMC inference on each simulation for 10^6^ iterations, logging every 10^3^ iteration, and discarding the first 2 × 10^5^ samples as burn-in. These guaranteed Effective Sample Sizes of at least 300 across all parameters. We used weakly informative priors for most parameters. Specifically, we used a uniform U(0, 100) for ln(λ_*r*_), ln(*µ*_*r*_) and *E*, an Inverse Gamma Γ^−1^(0.5, 0.1) for *σ*_λ_, a Gaussian N(0, 10) for *α*, and a Gamma Γ(1, 1) (*i*.*e*., Exp(1)) for *µ*. Initial simulations showed that for trees with less than 100 tips, *σ*_*µ*_ would largely follow the prior, which, given the high variance of our prior, would yield very bad mixing and numerical inaccuracies. Thus, we used the stronger prior for *σ*_*µ*_ of Γ^−1^(5, 0.5), which concentrates more density on parameter values closer to 0.

**Figure 2.**
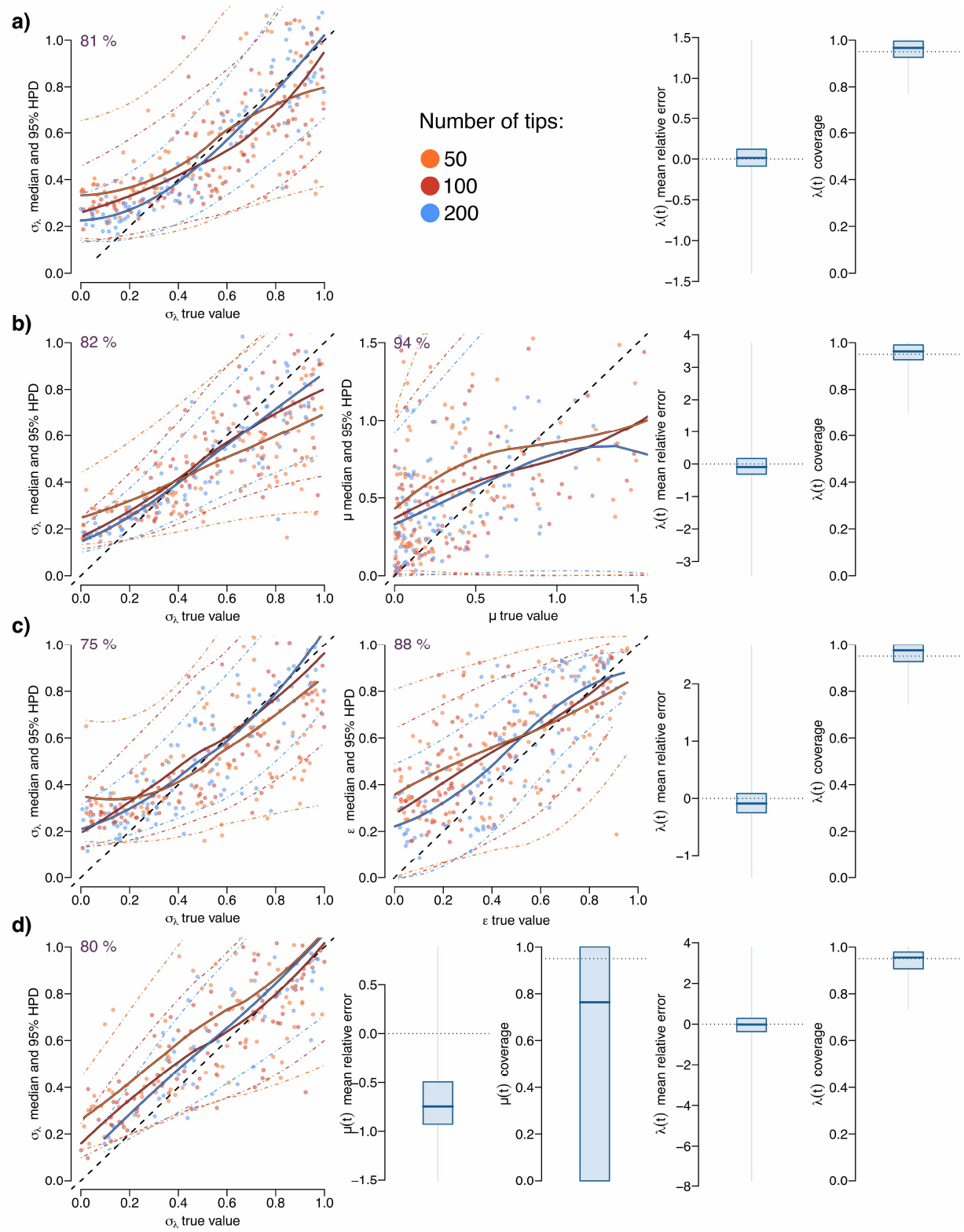
Validation of diffusion model without drift. Comparison between simulated and estimated values for **a)** pure-birth diffusion (‘PBD’: *µ*(*t*) = 0), **b)** constant-extinction diffusion (‘CED’: *µ*(*t*) = *µ*), **c)** constant-turnover diffusion (‘CTD’: *µ*(*t*) = *Eλ*(*t*)), and **d)** extinction-diffusion (‘BDD’: *µ*(*t*)), for 50, 100 and 200 tip simulations. For *σ*_λ_, *µ*, and *E*, we show the running means across simulations for the median estimates and the 95% HPD for each of the tree sizes, respectively, and the 1:1 line (black dashed). In the upper left corner we show the coverage for those parameters. For λ(*t*) (and, when applicable, *µ*(*t*)) we estimate the percentage average relative error across a series of fixed time points along the tree. Similarly, we calculate the coverage by counting the number of true values that fall within the 95% HPD in the estimates across those same fixed time points.

**Figure 3.**
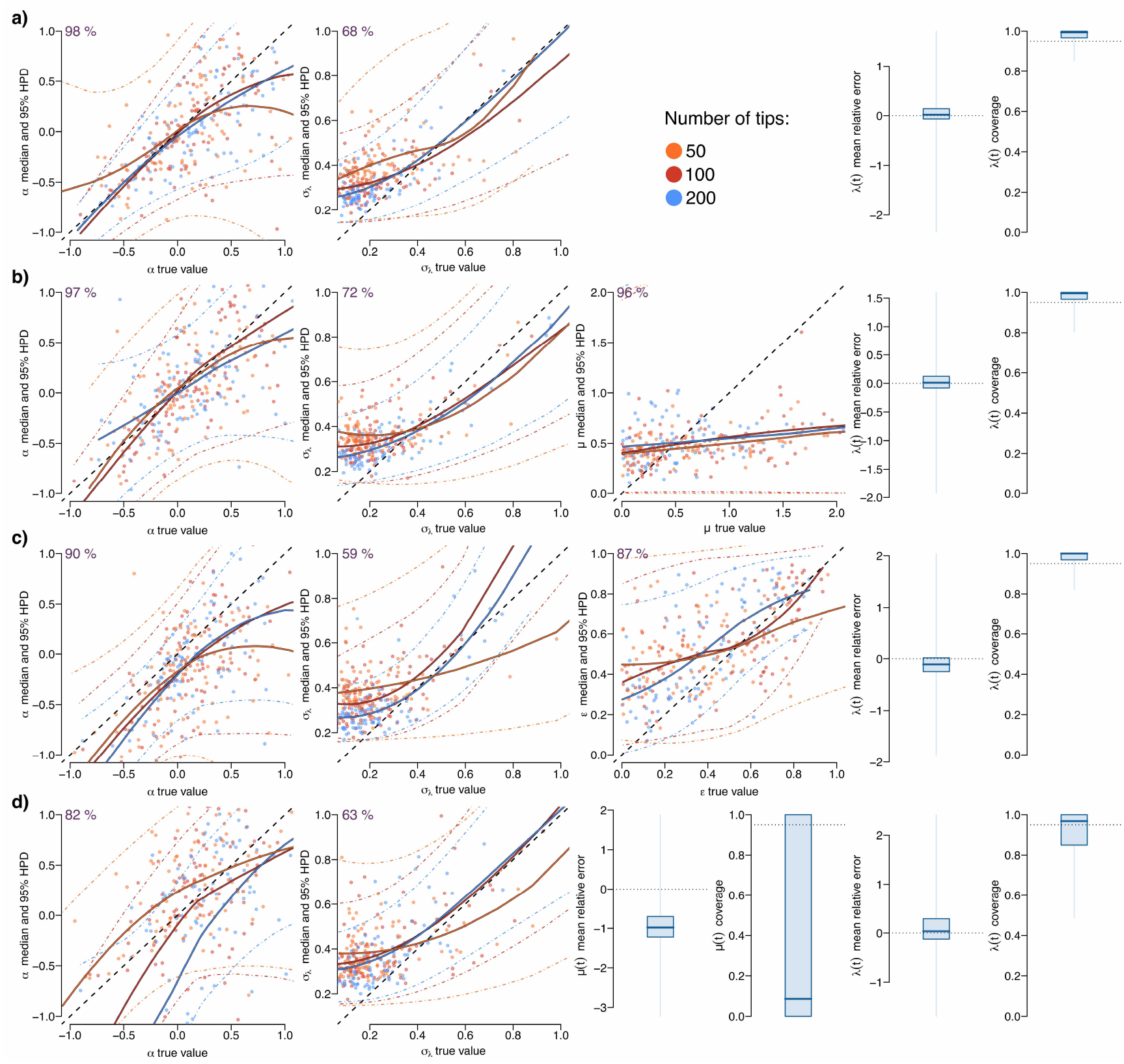
Validation of diffusion model with drift. Comparison between simulated and estimated values when including the drift parameter *α* for **a)** PBD, **b)** CED, **c)** CTD and **d)** BDD, for 50, 100 and 200 tip simulations. Plot details as in Figure 2.

#### Statistical coverage and accuracy

Figure 2 & 3 show the simulation results without and with the drift parameter *α*, respectively. Across all scenarios we have a median coverage of the speciation rates along each tree above *ca* 95% (a coverage of 95% for one simulation indicates that 95% of the true speciation rates across the phylogenetic tree are within the 95% Highest Posterior Density (HPD) intervals of the estimated speciation rates). Furthermore, the median average relative error (*i*.*e*., ln(λ_*true*_(*t*_*i*_))*/*ln(λ_*inferred*_(*t*_*i*_)) − 1)) for the diffusion of speciation rates is close to 0 across most simulations, supporting no strong bias in the model’s estimates (Fig 2 & 3). Nonetheless, for CTD we find a small downward bias (*i*.*e*., median of 0.2 relative log units) in speciation rates. Finally, the diffusion coefficient for speciation *σ*_λ_ shows adequate accuracy and coverage across all simulations (Fig 2 & 3), taking into account the influence of its prior for smaller trees.

As expected, the influence of the prior is proportional to the amount of data (*i*.*e*., tree size): Supplementary Figure 2 shows the influence of two different Inverse Gamma priors using the same data on the posterior distribution of *σ*_λ_ and compares coverage and accuracy across simulations with increasing tree size. Overall, *σ*_λ_ is overestimated in values close to 0 of parameter space (*i*.*e*., *σ*_λ_ < 0.1) when using an Inverse Gamma prior of Γ^−1^(0.5, 0.1). In this range of parameter space, using an Inverse Gamma prior with higher density towards 0 (*e*.*g*., Γ^−1^(0.5, 0.1)) can increase coverage, yet lead to some underestimation when rates are very high. While, as with all Bayesian analyses, the choice of prior is important and should be carefully considered, the underlying rate estimates do not suffer in statistical coverage nor accuracy, and the prior influence dwindles with increasing tree size (Fig S2).

Without drift, extinction rates for constant-extinction (*µ*(*t*) = *µ*) seem to be underestimated for trees with 50 tips, but gradually improve with trees of 100 and 200 tips; still, the 95% HPD coverage across these simulations is of 95% (Fig 2b). For constant turnover (*µ*(*t*) = *Eλ*(*t*)), *E* estimates are become more accurate and coverage is better as one increases tree size. For the BDD (*µ*(*t*)), extinction rates have very poor coverage and a underestimation bias, suggesting that, for extant-only small trees (at least to 200 tips), extinction diffusion dynamics seem not to be recoverable.

With drift, we get good estimates and coverage of the drift, *α*, and the diffusion, *σ*_λ_, parameters (Fig 3) across diffusion models (again taking into account the influence of the prior on the diffusion coefficient for small tree sizes; Fig S2). For CTD, we find good estimates of turnover rates, *E*, with good accuracy increasing with tree size. For CED, however, accuracy is low and, at least for the range of tree sizes used, does not seem to improve with more data (at least up to trees with 200 tips). HPD coverage of *µ* remains very high since the posterior distribution has high variance. For the BDD model, median extinction diffusion coverage is very low and shows a strong downward bias, again suggesting that either more data is needed or an impossibility to appropriately recover diffusing extinction rates. Note, however, this does not lead to non-identifiablity, but rather to biased estimates in extinction rates. While these results reinforce known difficulties of retrieving extinction rates from extant-only trees, particularly in our model with lineage specific speciation diffusion with drift, speciation rates continue displaying good statistical behavior.

#### Results summaries

On top of returning posterior samples for all hyperparameters, our model returns a posterior sample of data augmented trees, each with unobserved lineages that went extinct and their latent speciation, and in the case of BDD, extinction, rates. We summarize rate patterns in two ways. First, to provide posterior speciation and extinction (only in BDD) rate distributions across the tree, we remove all unobserved (data augmented) branches from the data augmented trees, and then estimate the posterior distribution for each λ_*i*_ in the reconstructed tree. Note that we can only estimate a posterior distribution of rates along the branches of the reconstructed tree, since this is the only part that remains fixed across the whole MCMC run. For clarity, we dub these estimates “posterior reconstructed rates”, which we can summarize at any point along the phylogenetic tree, or take their geometric mean across contemporary lineages through time.

Second, we estimate average rates through time taking into account all data augmented lineages. Here, we take each data augmented tree sample and estimate the cross-lineage geometric mean of their rates through time. We call these rates “posterior DA rates”.

#### Temporal patterns

In simulations of our diffusion models, the drift parameter controls whether the median speciation rates increase or decrease through time. Nonetheless, we wanted to further test the flexibility of our diffusion models in capturing temporal heterogeneity that is neither linear nor constant, making our models more useful when applied to empirical data. Thus, we performed a simulation scenario in which speciation rates were low, suddenly increased and then decreased, while maintaining branch heterogeneity, and with some extinction diffusion, as shown in Figure 4a. We then performed inference on this simulation using PBD (Fig 4b), CED (Fig 4c), CTD (Fig 4b), and BDD (Fig 4e,f). Remarkably, we find that all diffusion models are able to capture this temporal fluctuation in speciation rates, independent of our assumption on extinction rates.

**Figure 4.**
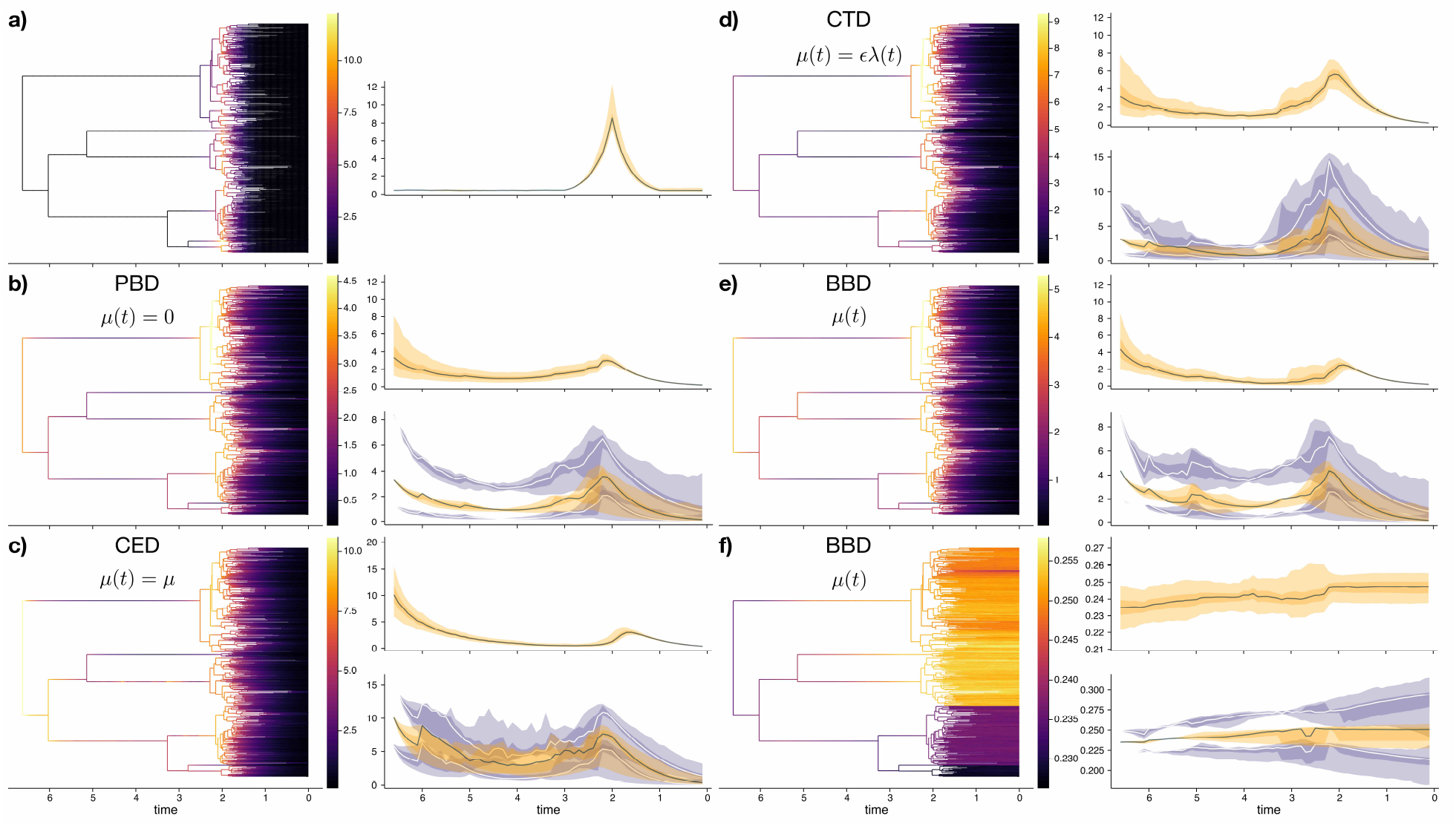
Common temporal patterns through time. **a)** Simulated tree (*left*) with dramatic temporal variation in speciation rates affecting all lineages simultaneously (*right* : geometric average λ(*t*) across lineages and 95% quantiles), small inter-lineage variance *σ*_λ_ = 0.1 and a constant extinction rate of 0.2. **b-f)** Show inferred patterns given the simulated tree. For each, *Left* : Median posterior reconstructed rates across the tree (speciation for ‘b-e’ and extinction for ‘f’); *Upper right* : posterior DA rates and 95% Credible Intervals (CI) across all data augmented trees; *Bottom right* : posterior median and 95% CI reconstructed rates. Inferred models are: **b)** PBD (*i*.*e*., *µ*(*t*) = 0), **c)** CED (*i*.*e*., *µ*(*t*) = *µ*), **d)** CTD (*i*.*e*., *µ*(*t*) = *Eλ*(*t*)), and BDD (*i*.*e*., *µ*(*t*)) **e)** speciation and **f)** extinction rates.

#### Implementation

We implemented all these diversification methods in the ‘Tapestree’ package for Julia (Bezanson et al., 2017), available at https://github.com/ignacioq/Tapestree.jl. The software documentation details the simulation, inference and data structures implemented in the package as well as its summary and plotting capabilities.

### Mammal Birth-Death Diffusion

We characterize the across-lineage temporal diversification of mammals using our new diffusion models. We used the recent time-calibrated phylogenetic tree integrating phylogenomic data from Álvarez-Carretero et al. (2021), which comprises 4705 genetically represented extant lineages. Because many of these tips are still considered subspecies, we used only recognized species as tips for taxonomic consistency in the patterns observed, yielding a tree with 4071 species. We estimated clade-specific sampling fractions for each of 16 subclades we identified and which broadly correspond to those divisions used to infer the full mammal phylogenetic tree in Álvarez-Carretero et al. (2021). For this, we matched current mammals diversity following the Mammal Diversity Database (‘MDD’; 2022), which at the time of the analyses (May 2022) recognized 6457 total extant mammal species. We matched the species with the taxonomy in the MDD and estimated how many species were not represented in the tree to estimate species specific sampling fractions to be passed to the diffusion models (subclade specific sampling fractions are shown in Table S1). We ran 3 MCMC chains on the Maximum Clade Credibility (MCC) mammal phylogeny for 10^6^ iterations, saving every 10^3^ and discarding an additional 3 × 10^5^ as burn-in, for each of the diversification models: PBD, CED, CTD and BDD, both with and without the drift parameter. Finally, we added a constant turnover diffusion model fixing turnover to 1 (equal rates of speciation and extinction at any given time) following empirical paleontological evidence (Marshall, 2017).

As expected, we find substantial variation across surviving mammals lineages in speciation and extinction rates (Fig 5 & 6), with a median posterior diffusion coefficient for speciation rates of *σ*_λ_ = 0.117 spp/My^3/2^ and of extinction rates of *σ*_*µ*_ = 0.067 spp/My^3/2^. Indeed, median posterior reconstructed rates of speciation range from about 0.01 spp/My and of extinction of up to about 0.025 spp/My. We find that posterior reconstructed and DA speciation rates were stable or even slightly decreasing since the origin of extant mammals throughout the Jurassic and most of the Cretaceous, but that, during the late Cretaceous (*ca*. 80 Mya), there was a dramatic increase in speciation rates (Fig 5a,c). Thereafter, speciation rates remained stable until recently where a final surge in speciation rates is evinced, mostly driven by the recent fast diversification of rodents. These patterns of speciation rates were congruent across diffusion models, *i*.*e*., PBD (Fig S3), CED (Fig S4), CTD (Fig S5) and CTD with turnover fixed to 1 (Fig S6). Finally, diversity curves show different patterns across the different diffusion models (Fig 5 & Fig S3-S5), yet, regardless of the assumption in extinction, we find an initial slow accumulation of diversity with a sharp increase in the Late Cretaceous mirroring the temporal pattern of speciation rates.

**Figure 5.**
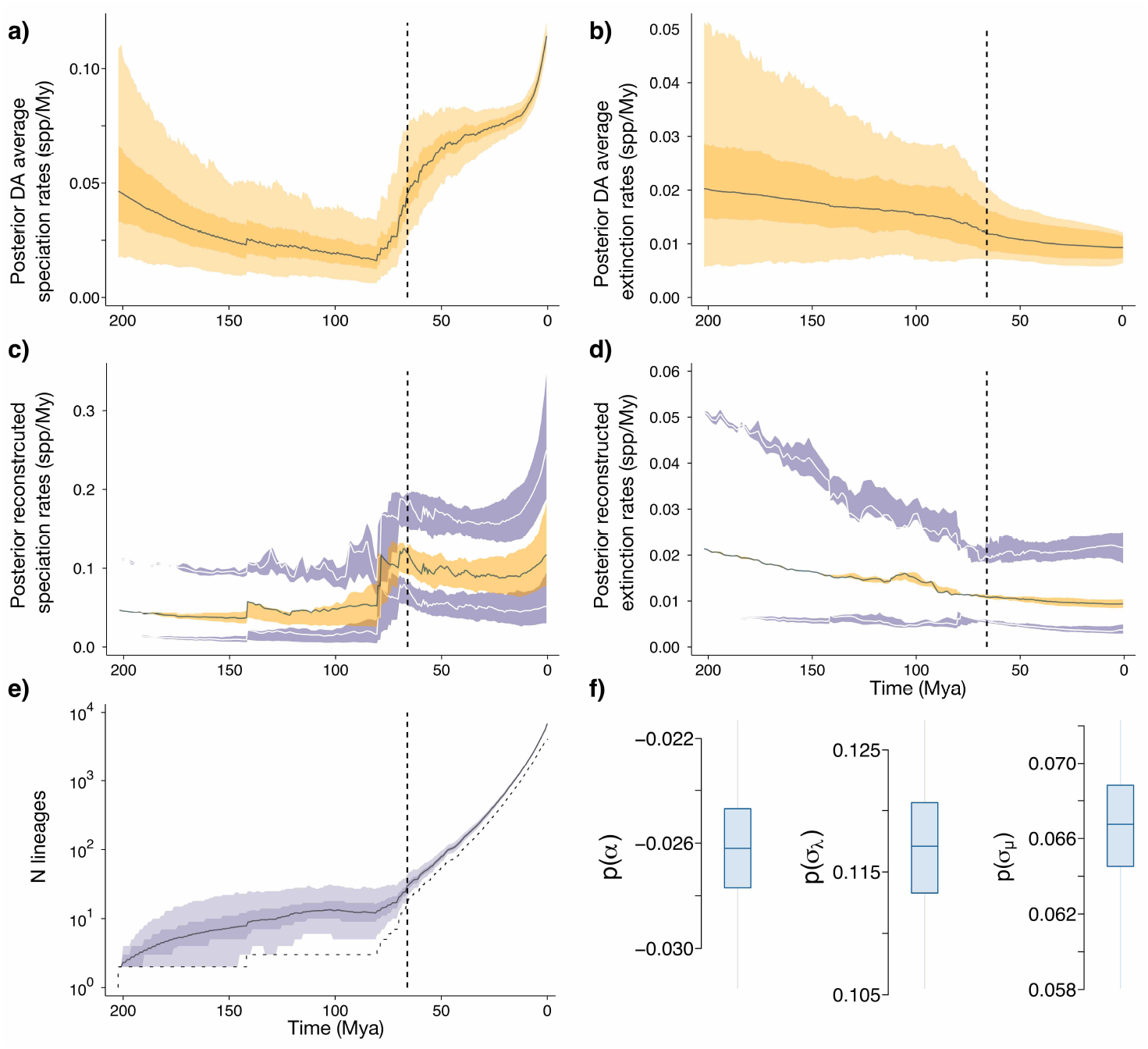
Birth-Death Diffusion for surviving mammals: overarching patterns. **a)** Posterior DA average speciation rates and **b)** extinction rates across all lineages across augmented tree samples. Solid line shows the median, darker orange the 50% CI, lighter orange the 95% CI across all data augmented trees. **c)** Posterior reconstructed speciation rates and **d)** extinction rates for median posterior tree (orange shading showing median and 50% quantiles across the lineages’ median posterior rates), and for 2.5% and 97.5% CI trees (purple shading showing median and 50% quantiles across the lineages’ 2.5% and 97.5% posterior rates, respectively). **e)** Distribution through time of the number of mammals lineages in the data augmented trees (solid line shows the median, darker purple the 50% CI, lighter purple the 95% CI). The lower dotted line shows the lineages in the reconstructed tree only. **f)** Posterior distributions (95 %, 50 % and median HPD) for BDD hyperparameters: drift *α*, diffusion in speciation rates *σ*_λ_ and in extinction rates *σ*_*µ*_.

**Figure 6.**
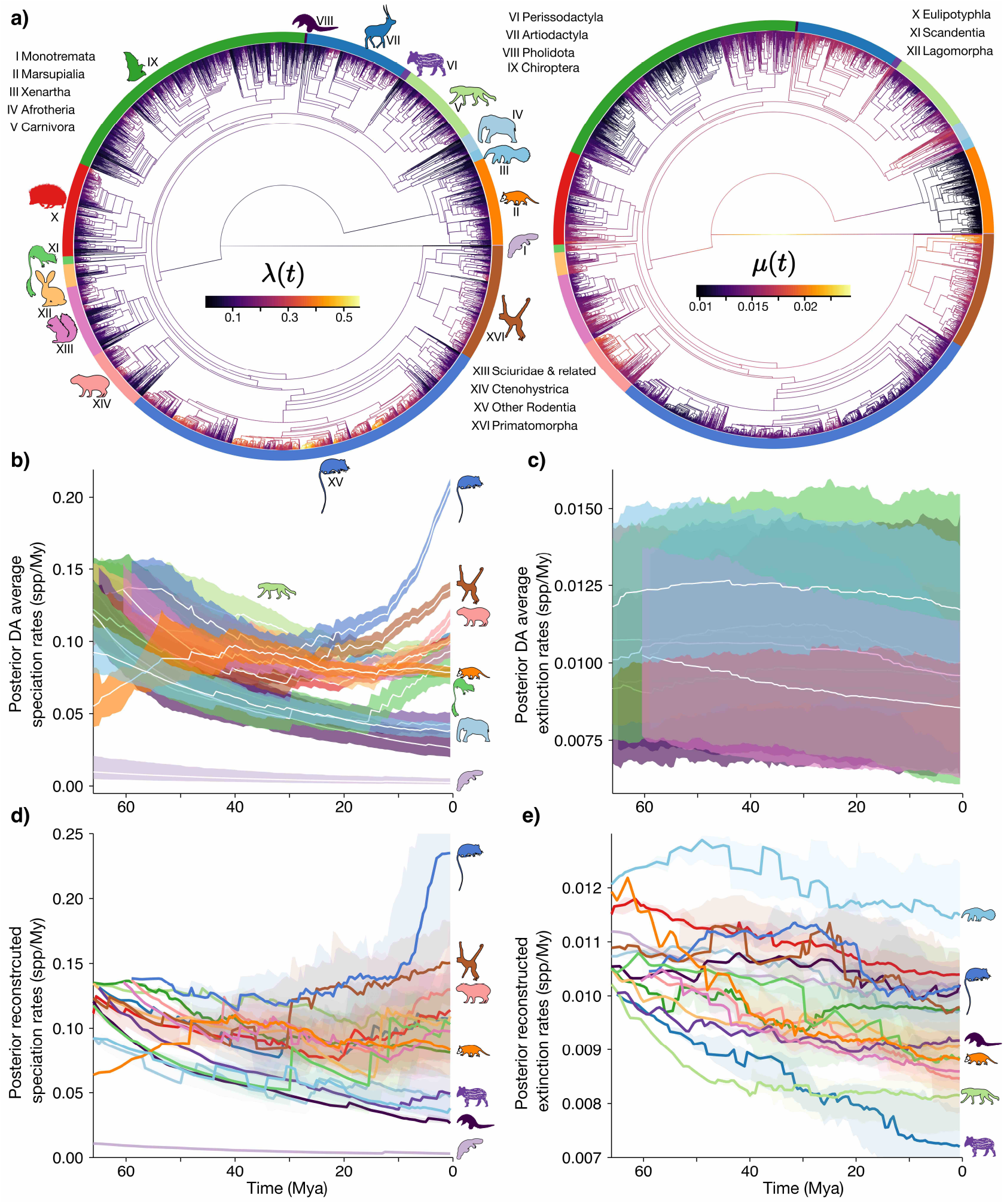
Birth-Death Diffusion for surviving mammals: lineage heterogeneity. **a)** Median posterior reconstructed speciation rates λ(*t*) and extinction rates *µ*(*t*) across the reconstructed mammals tree according to the Birth-Death Diffusion model (note that this are the same model results of Fig 5). Surrounding colors identify 16 mammal clades with en embedded species silhouette and roman numerals for identification. **b)** Posterior DA average speciation rates and **c)** extinction rates the 16 major clades highlighted in ‘a)’. Solid white line shows the median, while the darker shade the 50% CI across all data augmented trees. Silhouettes identify some focal clade patters. **d)** Posterior reconstructed speciation rates and **e)** extinction rates for median posterior tree (solid lines shows the median and light shading shows the 50% quantiles across the lineages’ median posterior rates).

## Discussion

We developed and implemented new flexible birth-death diffusion models, which allow fast estimation of speciation and extinction rates that vary instantaneously and continuously in time and across lineages. Our model provides increased resolution than other approaches assuming a few constant regime shifts across the tree (Rabosky, 2014; Höhna et al., 2019; Barido-Sottani et al., 2020; Vasconcelos et al., 2022), and even ‘ClaDS’, which assumes constancy in diversification rates within a branch (Maliet et al., 2019). Indeed, for any lineage, for any instantaneous point in time, our model returns a posterior distribution of speciation (and extinction) rates. Furthermore, we present four different models with different assumptions about extinction, three of which are analogous to ClaDS (*i*.*e*., ‘pure-birth’, ‘constant-extinction’ and ‘constant-turnover’), and a more flexible one that assumes extinction to also vary according to a GBM, the ‘BDD’ model. As with the data augmented implementation of ClaDS (Maliet and Morlon, 2021), our inference yields a set of data augmented posterior trees, that is, probable histories given the model and data, which can be used to estimate diversity through time and average rates (*e*.*g*., as we did for mammals in Fig 5 & 6), avoiding biases for not taking into account the dynamics of unobserved lineages on these estimates. We also generalize the data augmentation scheme for phylogenetic birth-death processes and show that it can be applied to other models of diversification for which the likelihood has no analytical solution or its calculation is computationally costly.

Our diffusion models assume that diversification rate variation is heritable, accumulates in a gradual fashion and in proportion to time. This is consistent with the idea that the interdependence of species traits with their environment at any given moment in time will largely govern their diversification rates. Heritable traits such as body size, dispersal and generation time, among others, are posited to influence rates of speciation and extinction, and result from a gradual, heritable evolutionary process (Heard, 1996). Further evidence of the heritability of diversification rates is found in the observed imbalance across empirical phylogenetic trees (Heard, 1996), as expected from speciation and extinction rates that evolve through time and are transferred into daughter species. However, non-heritable variation, such as environmental oscillations, are also thought of influencing diversification rates (Heard, 1996; Barnosky, 2001; Benton, 2009). We show that, while not explicitly accounted for, our diffusion models are able to detect overarching temporal trends while accounting for fine-grained lineage heterogeneity.

All diffusion models support the ‘early-rise’ hypothesis for the clades that led to the present-day richness of mammals (Bininda-Emonds et al., 2007; Meredith et al., 2011; Springer et al., 2019; Grossnickle et al., 2019). Specifically, we find that the origin of the major clades leading to extant mammal diversity results from a sharp average increase across contemporary lineages in speciation rates before the K-Pg extinction event (Fig 5). This refutes the explosive-rise hypothesis wherein mammals were suppressed in their ecomorphological and taxonomic diversity during the Cretaceous and experienced a release after the K-Pg extinction event (Archibald and Deutschman, 2001). Our results are concordant with findings using previous phylogenetic evidence from (Meredith et al., 2011), and are supported by other lines of recent paleontological evidence showing that a major ecomorphological diversification in mammal lineages following the rise of angiosperms occurred 10-20 My before the K-Pg (Wilson et al., 2012; Grossnickle and Newham, 2016; Halliday and Goswami, 2016; Grossnickle et al., 2019; Upham et al., 2021). Specifically, fossil dental and body size analyses reveal an expansion into herbivorous and carnivorous diets and increased body size disparity concomitant with taxonomic diversification (Grossnickle and Newham, 2016; Wilson, 2014).

Before the diversification burst of the Late Cretaceous, our diffusion models that include extinction exhibits low speciation rates coupled with high extinction rates rates, leading to a slow initial accumulation of mammal diversity (Fig 5c-g). This pattern is also mirrored in paleontological evidence from the Cretaceous Terrestrial Revolution that resulted in substantial lineage turnover (Luo, 2007; Grossnickle and Polly, 2013; Benson et al., 2013, 2016; Grossnickle et al., 2019). While we do not find support for a delayed-rise only scenario of mammals, diversification rates remained high after the K-Pg during most of the Paleogene with a sustained increase in diversity (Fig 5c-g). This findings contrast with the results from (Meredith et al., 2011), wherein speciation rates increase at *ca*. 83 Mya but decrease sharply again at *ca*. 78 Mya, and agrees with paleontological evidence of continued ecological and taxonomic diversification (O’Leary et al., 2013; Grossnickle et al., 2019). Our results also show no effect of the K-Pg extinction event on diversification rates of the surviving mammals lineages but instead show a continued acceleration until the Eocene, concomitant with the radiation of many of the crown group members of extant placental orders (Grossnickle et al., 2019). Regardless of the assumption in extinction rates (*i*.*e*., no extinction, constant extinction, constant turnover, constant turnover of 1 or extinction following a diffusion process), our results suggest that the K-Pg did not drive the explosive radiation of present-day mammals (Fig 5,S3-S6). Instead, while the accelerated diversification of surviving mammals started before the K-Pg, the aftermath of this extinction event allowed most lineages to maintain comparable (or even higher) levels of high rates of diversification up to the present.

Underlying these overarching processes, our model reveals substantial lineage and time heterogeneity of diversification rates across the mammal tree (Fig 6). To illustrate, we find posterior median lineage speciation rates that range from almost 0 spp/My in Monotremes, up to more than 0.2 spp/My in some Rodents. Concomitant to their high extant diversity, mouse-related Rodents (excluding Ctenohystrica and Sciuridae and related clades) exhibit a dramatic surge in their speciation rates from *ca*. the start of the Miocene (Fig 6). Multiple hypotheses, such as developing hypsodonty, colonization of South America, environmental changes and extinction of competitors, have been proposed to explain this evolutionary radiation (Fabre et al., 2012). Speciation rates in Primates also follow a post-Oligocene increase, yet this was preceded by a sharp decrease after the K-Pg up to the Oligocene, coinciding with a terrestrial fauna turnover event from glaciation and cooling (Springer et al., 2012). For their part, Marsupials show an increase of speciation rates during the Paleocene, correlated with biogeographic dispersal from South America to Australia through Antarctica (Nilsson et al., 2004), but then remained stable. These examples of clade specific routes to extant diversity demonstrate the strength and flexibility of our birth-death diffusion models in capturing how lineage-specific diversification rates evolve in continuous time.

Our model and results assume that the phylogenetic tree used is the ‘true’ tree of extant mammalian evolution, and relies on its specific topology and timing of speciation events. Consequently, we use the most up-to-date phylogenetic time-tree of mammals, built with a novel Bayesian framework that incorporated 72 mammalian genomes and thousands of species genetic data in an integrated framework, with substantial less uncertainty in the timing of past cladogenetic events (Álvarez-Carretero et al., 2021). Our diversification results contrast with previous phylogenetic analyses that used different model assumptions and phylogenetic trees. Bininda-Emonds et al. (2007) developed the first species-level phylogenetic tree and estimated changes in the slope in the Lineage-Through-Time (LTT) plot to show a diversification peak at around 90 Mya, followed by a decrease until the K-Pg, where rates increased again until the very present. Using the same phylogenetic tree, Stadler (2011a) demonstrated that using the LTT slope as means to estimate temporal changes in diversification rates can lead to biased results, and developed a birth-death model where contemporary lineages experience the same diversification rates (*i*.*e*., ‘lineage-homogeneous’), but are allowed to shift at different epochs. Using this model, a peak in mammalian diversification rates around 33 Mya was inferred, followed by stable high rates until near the present, wherein they declined. Meredith et al. (2011) built a family level tree with 169 mammal species and recovered a diversification peak that lasted only during *ca*. 83-78 Mya. Using the same model, but with an updated tree and dating scheme of placental mammals, Liu et al. (2017) found an upward shift around 90 Mya followed by a downward shift around 52 Mya, and no effect of the K-Pg. Finally, using the tree from Upham et al. (2019), Upham et al. (2021) performed temporal diversification analyses under a model of lineage-heterogeneous rates resulting from discrete shifts (Rabosky, 2014) (but see Moore et al. (2016) for concerns on this method) and found a burst in speciation rates in the Late Cretaceous followed by a steady increase towards the present.

A more general caveat persists for diversification analyses that are conducted on fixed phylogenetic trees (the great majority), which, in turn, were usually inferred using a dating method that does entail explicit (typically through constant pure-birth or birth-death priors) or implicit assumptions about the underlying diversification process. Ultimately, however, a joint model where co-estimation of divergence times and diversification rates would be more appropriate, and has recently been developed for some birth-death models with discrete shifts (Kühnert et al., 2016; Höhna et al., 2016; Barido-Sottani et al., 2020). The data augmented approaches developed here could be used as priors over divergence times in an integrative approach and is an exciting avenue for future research.

As with most diversification models based on extant-taxa alone, our model and results are susceptible to inferential limits, specially as one moves into the deep past. For instance, Louca and Pennell (2020) showed that for time-varying lineage-homogeneous speciation and extinction rates and in the absence of any constrain on the functional form of the time-varying rate functions, there are an infinite number of such functions that return the same likelihood for any extant-only time-tree. Our model overcomes this particular non-identifiability issue by incorporating lineage-heterogeneous rates informed by topology by means of speciation and extinction rates following a GBM diffusion model. More applicable, however, are concerns in reliably estimating extinction rates from extant-only time-trees (Rabosky, 2010; Beaulieu and O’Meara, 2015; Louca and Pennell, 2021). Note that our inference model is similar, yet more complex, to the simulation model in Beaulieu and O’Meara (2015): they simulated trees with branch-constant rates that are inherited with a change proportional to the product of a diffusion rate and the squared root of elapsed time, rectifying the time proportionality of the diffusion rates used by Rabosky (2010), who used the product of time (rather than squared root of time) with the diffusion rate. Using these simulations, Rabosky (2010) and Beaulieu and O’Meara (2015) found that a constant rate birth-death model will overestimate extinction and lead to spurious results if the variance in speciation rates was high. Our model explicitly takes into account this type of heritability and thus is immune to this specific across-lineage heterogeneity that could lead to inaccurate inferences when assuming a simpler model.

This is not to say that other sources of variation in diversification could not potentially bias inference. From the different extinction assumptions in our diffusion models, the best behavior is achieved for constant-turnover (CTD) and constant-extinction (CED), with the extinction diffusion model (BDD) only achieving fair accuracy with sufficient data. Nonetheless, we are cautious on over-interpreting what these extinction estimates represent, particularly when faced with empirical data. We emphasize that our results reflect the diversification dynamics of those mammal lineages under a still simple model of diversification, and, using extant-species alone trees could prove insufficient in recovering very high extinction rates in the past (Marshall, 2017). For instance, without fossil information the probability of a mammal clade that diversified greatly yet went extinct, such as the Cimolodontan multituberculates (Grossnickle et al., 2019), is marginal under birth-death models conditioned on surviving taxa and thus is unlikely to be inferred as the main diversification history. Furthermore, our model does not account for episodic mass extinction events, which could substantially enhance the biological realism of our diversification histories, conditional on having a good estimate of the proportion of lineages that went extinct. Our model and implementation renders the future incorporation of fossils into the model straightforward and opens an exciting avenue for future macroevolutionary research as phylogenetic trees that combine neontological and paleontological data become increasingly available.

One of the main properties of the diffusion models is that speciation (and extinction in the BDD) vary continuously and stochastically within any lineage at any moment in time. On one hand, this unlocks a new series of possibilities for answering questions about evolutionary radiations. Specifically, it provides the foundation in which to test drivers of diversification rates that also vary instantaneously through time and across lineages, such as concomitant evolution of species intrinsic traits or abiotic factors, including environmental fluctuations, while accounting for independent, stochastic variation. This would be the continuous counterpart to the hidden states framework that considers residual variation in State dependent Speciation and Extinction methods (Beaulieu and O’Meara, 2016), which have proved pivotal in assessing discrete trait drivers of diversification. ‘QuaSSE’ provides a correlation test between diversification and a continuous trait FitzJohn (2010), but, aside of not accounting for residual variation (increasing Type I error), uses a numerical method that would not scale-up well with the number of Brownian processes. Conversely, we have shown with our data augmented framework that this would not be a major limiting factor. More generally, our framework is analogous to a simple linear regression in having both a deterministic effect (*i*.*e*., slope) together with unexplained variance. This analogy illustrates the strength of such an approach for it will be contentious to perform inference using only a linear slope to explain the association between two variables. Future advancements in this direction will help in identifying the underlying processes governing macroevolutionary dynamics.

With our new model, there are now three different paradigms for modeling lineage-specific variation in diversification rates: rare discrete anagenetic shifts (*e*.*g*., ‘BAMM’, ‘MiSSE’, and related; Rabosky, 2014; Höhna et al., 2019; Vasconcelos et al., 2022), discrete cladogenetic shifts (*i*.*e*., ‘ClaDS’; Maliet et al., 2019), and, with our models, continuous anagenetic diffusion. The level at which diversification rates change in a few discrete bouts or gradually is unknown, and probably both types of events have contributed in generating diversification rate variance (Barnosky, 2001; Benton, 2009; Maliet et al., 2019). Arguably, even those novel traits thought of substantially impacting lineage diversification rates (*e*.*g*., ‘key’ innovations) do not appear ‘instantaneously’ in time, but take some time to evolve, even if brief (Mayr, 1963; Heard and Hauser, 1995; Hunter, 1998). Similarly, it remains an open and intriguing question whether changes in diversification rates occur mostly at cladogenesis, a likely prediction under punctuated equilibrium with trait dependence of diversification rates (Gould and Eldredge, 1972; Maliet et al., 2019), or evolve throughout a lineage’s lifetime. We are only aware of one phylogenetic study that compared BAMM, a model of few discrete shifts (Rabosky, 2014), with ClaDS, a model with many branch specific shifts (Maliet et al., 2019), and found higher Bayesian support for the latter across some empirical trees, supporting the idea that most variation in diversification rates are accumulated through frequent small-scale variation (Ronquist et al., 2021). Given that all these processes could be present to some degree in empirical datasets, comparing models with exclusive processes may not be particularly meaningful. A promising and exciting perspective of the data augmentation methods introduced here is that they can easily be recruited to develop models that would combine those hitherto disconnected processes into a joint model, where their respective contributions could be directly estimated.

Our new model development uncovered a clear signal of an early-rise of modern mammals, refuting the idea that suppression before the K-Pg was limiting the diversification of those mammal lineages that survived to the present, but rather suggest the angiosperm radiation opened substantial ecological opportunities for these lineages to diversify in the Late Cretaceous and up to the present. These results derive from inferring rates of speciation and extinction that can vary at any time for any lineage, as theoretically expected from the interplay of intrinsic and extrinsic factors acting at a given time on species, and opens new modeling and computational possibilities in the study of macroevolutionary dynamics.

## Acknowledgments and funding

This project has received funding from the European Union’s Horizon 2020 research and innovation programme under the Marie Sklodowska-Curie grant agreement No 897225 for IQ.

